# Probing the toxic interactions between the reactive dye Drimaren Red and Human Serum Albumin

**DOI:** 10.1101/2021.07.17.452798

**Authors:** Thaís Meira Menezes, Caio Rodrigo Dias de Assis, Antônio Marinho da Silva Neto, Priscila Gubert, Marcos Gomes Ghislandi, Jorge Luiz Neves

## Abstract

Azo dyes like Drimaren Red CL-5B (DR, CI Reactive Red 241) represent a class of compounds extensively used in the textile industry and are extremely dangerous to the environment and human health. Therefore, understanding the binding characteristics between such substances and biological macromolecules is essential from a toxic-kinetic perspective. The molecular interaction between DR and Human Serum Albumin (HSA) was investigated through spectroscopic techniques and molecular docking approaches. The results indicate that DR quenches HSA fluorescence following a static mechanism (corroborated by UV-Vis studies) with a moderate interaction (K_a_~10^5^ M^−1^), guided by electrostatic interactions (ΔS°> 0 and ΔH°< 0). DR is 5.52 nm distant from fluorophore residue Trp-214 (according to FRET investigations), and the interaction is mainly related to Tyr residues (as revealed by synchronous fluorescence). The Ellman assay identified a decrease in the content of HSA free thiol. The results of the RLS demonstrate that there are HSA alterations, suggesting damage to the confirmation of the protein. Molecular docking suggests the binding site of DR was located in subdomain IIB HSA, corroborating the experimental properties. Finally, the results suggest a high potential for DR toxicity triggered by contact with key proteins, which affects the biomolecule functionalities.

## 1. INTRODUCTION

The textile industry comprises a complex network of processes that constantly exacerbate the use of organic dyes (Haji and Naebe, 2020), which are often improperly released into the environment (Gita et al., 2017; Zafiu et al., 2021) and causes recurrent human health problems (Ding et al., 2011; Kooravand et al., 2021; Naveenraj et al., 2018; Rovira and Domingo, 2019; Tkaczyk et al., 2020; Zhang et al., 2009). In this regard, most textile dyes are potentially toxic, carcinogenic, mutagenic and teratogenic (Carneiro et al., 2010; Tan et al., 2020; Yek et al., 2020). Furthermore, improper dye residue disposal causes soil and groundwater contamination, reducing the water quality for humans and aquatic ecosystems. Moreover, dye-contaminated water in crop irrigation systems can directly affect the entire food chain (Kishor et al., 2021). Therefore, as a hazardous material, their water body accumulation has been a concern across the globe.

Reactive dyes are substances containing electrophilic groups in their structure. They can react with nucleophilic groups present on the surface of fabrics and fibers (OH, NH and SH groups), forming covalent bonds (Benkhaya et al., 2020). The desired properties for the application in the textile industry are related to their chromophore and the reactive (H. Zhang et al., 2018) groups. However, considering the chemical particularities of these substances and complex biological systems, the contact of reactive dyes with biomacromolecules (such as proteins, which have in their structure a large number of nucleophilic groups (Mu◻ller, 2018)) can result in alterations and loss of molecular physiological functions (LoPachin and DeCaprio, 2005). In addition, the electrophilic group-containing organic compounds (present in drinking water) have been reported to induce toxicity and be mediators of the carcinogenesis process (Prasse, 2021). The mentioned effects can arise from a series of biochemical processes that start from non-covalent interactions between the biomacromolecule and the dye, forming toxic adducts for the organism (Allen et al., 2014). For example, Drimaren Red CL-5B (CI Reactive Red 241, DR) is a reactive dye extensively used in the textile and leather industry (Javeda et al., 2019; Nisar et al., 2017; Sajjad et al., 2021; Santra et al., 2020; Soria-Sánchez et al., 2012) and exhibits in its structure (Fig. 1), the monochlorotriazine and the sulfatoethylsulfonyl reactive groups (Shao et al., 2014). In addition, the azo and anthraquinone groups act as chromophoric agents (Zhang et al., 2019), while the negative charged group SO^3-^ provides high solubility. However, investigations concerning the DR impact in biological systems lack in the scientific literature.

**Fig. 1.**
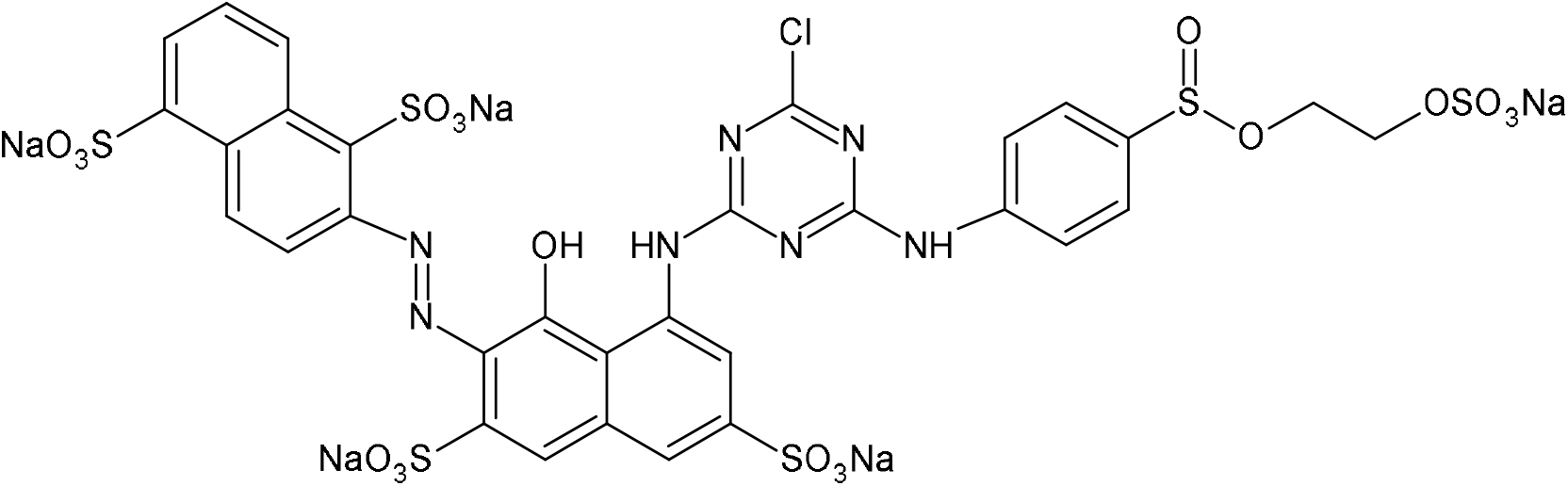
Structure of drimaren red CL-5B dye.

At the organism level, the effects caused by xenobiotic compounds are commonly mediated by carrier proteins present in blood plasma (Manjubaashini et al., 2018). For example, human serum albumin (HSA), a major plasma protein, carries out the transport of both endogenous compounds (fatty acids, hormones) and exogenous compounds (drugs, toxic substances) by the circulatory system (Maciążek-Jurczyk et al., 2018; Yamasaki et al., 2013), among several functions. HSA has 585 amino acid residues, having only one tryptophan residue (Trp-214) in its entire structure (Birkett et al., 1977). This protein has 35 cysteine residues, 1 of which has the free SH group (Cys-34) (Quinlan et al., 2005), and 34 make disulfide bridges that contribute to the stability of its tertiary structure (Paris et al., 2012). Their binding sites are three homologous domains, I, II and III, divided into subdomains A and B (Sugio et al., 1999). HSA has three main binding sites, located in subdomain IIA (Sudlow Site I), subdomain IIIA (Sudlow Site II) and subdomain IB (site III) (Sudlow et al., 1976; Zsila, 2013). The thiol group in Cys-34 also represents an important binding site located in subdomain IA, given its performance in the mechanisms of redox reactions (Watanabe et al., 2017). The HSA demonstrates a remarkable ability to form complexes with xenobiotics, directly influencing the toxicokinetics of hazardous substances (Faisal et al., 2020).

Here, the DR-HSA interaction and the protein alterations (induced by DR under physiological conditions) have been investigated, using multi-spectroscopic and molecular docking tools to clarify the potential toxicity of DR at the biomolecular level.

## 2. MATERIALS AND METHODS

### 2.1. Materials

Human serum albumin (fraction V) 98% and 5,5’-dithiobis-2-nitrobenzoic acid (DTNB) were purchased from Sigma Aldrich (USA) and used without any prior purification. Drimaren Red CL-5B (CI 18220) was purchased from Clariant SA. The HSA stock solution was prepared at a concentration of 5 μM using a buffer solution containing 0.5 M Tris-HCl (pH 7.4). The DR stock solution was prepared by dissolving the compound in Tris-HCl buffer at a concentration of 8.8 x10^−4^ M.

### 2.2. Methods

#### 2.2.1. Fluorescence quenching measures

The fluorescence quenching studies were measured on a Spectrofluorimeter Model F-7100 (Hitachi, Japan) coupled to a thermostated bath. The spectra were measured with excitation at 280 nm and excitation/emission slit width of 5 nm. The emission spectra were recorded with a scanning interval between 290 and 400 nm using a quartz cuvette. An HSA solution (0.6 μM) was used, and the DR was added to the solution in increasing concentrations between 0 and 17.60 μM. The experiments were carried out on 296, 303 and 310 K, being the internal filter effect corrected using equation (Musa et al., 2020):

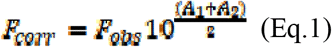

F_corr_ is the corrected, and F_obs_ is the observed fluorescence intensity. A_1_ and A_2_ are the DR absorbances at the excitation and emission wavelength, respectively. The Stern-Volmer equation (Ali et al., 2020):

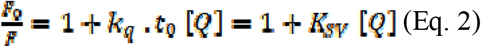

Moreover, the modified Stern-Volmer equation was used to calculate the parameters that demonstrate the association constant:

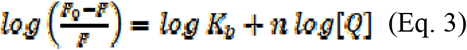

F and F_0_ are the fluorescence intensity at the maximum emission wavelength in the presence and absence of the suppressor, respectively. [Q] is the fluorescence suppressor concentration, kq is the bimolecular quenching constant, t_0_ is the average biomolecule lifetime in the absence of the quencher (6.38 x 10^−9^ s) (Abou-Zied and Al-Shihi, 2008), n is the stoichiometry of the binding and K_b_ is the binding constant. At varying temperatures, the equation

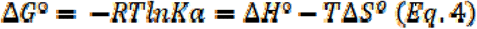

provides the values of enthalpy (ΔH°), entropy (ΔS°) and Gibbs-free energy (ΔG°), where R is the universal gas constant (R = 8,314 J.mol^−1^.K^−1^), and T is the temperature used during the experiments

#### 2.2.2. Measuring Forster Resonance Energy Transfer (FRET)

For the FRET study, the DR absorption, HSA emission and HSA-DR emission spectra were measured in 1: 1 stoichiometry. The binding distance (r) between the donor (Trp-214 of HSA) and the acceptor (DR) was evaluated according to Foerster’s resonance energy transfer theory (Moeinpour et al., 2016). According to Foerster’s theory, r can be measured from the equation 5:

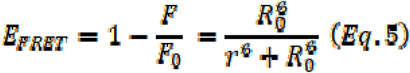

Where R_0_ is the Foerster distance at which the FRET efficiency reaches a value of 50%. The value of R_0_ is obtained by the equation:

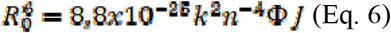

Furthermore, the value of J is obtained by:

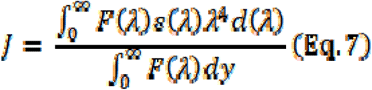

Where k^2^ is the spatial orientation factor of the dipole, n is the refractive index of the medium, Ф is the quantum yield of the donor’s fluorescence J is the donor’s overlapping integral’s fluorescence emission spectrum with the acceptor’s absorption spectrum. In this study, it is assumed that k^2^ = 2/3 n = 1.336, and Ф = 0.15 (Naik and Jaldappagari, 2019).

#### 2.2.3. Synchronous Fluorescence Measurements

The HSA synchronous fluorescence experiments at 0.6 μM were recorded on the Jasco spectrofluorimeter, model FP-6300 (Kyoto, Japan) in the absence and presence of varying concentrations of DR (0 to 17.60 μM), simultaneously carrying out the variation of the excitation monochromators and emission (Δλ = λ_em_−λ_ex_), with Δλ = 15 nm for Tyr residues and Δλ = 60 nm for Trp residues. In addition, the synchronous fluorescence quenching ratio (RSFQ%) was measured using equation (Zhao et al., 2020):

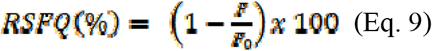

#### 2.2.4. UV-Vis Absorption Spectroscopy

The UV-Vis absorption spectra were measured on a spectrophotometer model Lambda 650 (Perkin Elmer, USA) using a cuvette of 1 cm path length, ranging from 200 nm to 400 nm at room temperature. DR aliquots (0-8.8 μM) were added to an HSA solution (0.6 μM) to the system at concentrations of 0 and 8.8 μM. The absorption spectrum of DR solutions in the absence of the protein was also recorded for correction of fluorescence inner filter.

#### 2.2.5. Ellman Assay

The content of reactive thiol groups present in protein samples was quantified using the modified Ellman method (Pavićević et al., 2014). HSA in the absence and presence of variated concentrations of DR were diluted in 0.5 M Tris–HCl (pH 7.4) and incubated with DTNB (100 μM) and EDTA (10 mM) for 15 minutes at room temperature. The absorbance of the reactional mixture was recorded at 412 nm. For the calculation of the thiol content, the molar extinction coefficient of 14,150 mol^−1^.L.cm^−1^ was used.

#### 2.2.6. Rayleigh light scattering (RLS) measurements

Rayleigh light scattering measurements were recorded using a Jasco spectrofluorometer, model FP-6300 (Kyoto, Japan), at room temperature in a 1 cm path length cuvette. The spectra were measured in synchronous mode, where Δλ = 0 nm in the region between 200 and 600 nm. The excitation and emission slit widths were both set at 1.5 nm.

#### 2.2.7. Molecular Docking

Molecular docking studies were conducted to investigate plausible structural factors which could help to explain the experimental observations obtained. First, the SMILE molecular representation of Drimaren Red chemical information was obtained by using open Babel. Then, it was used to obtain 3D structure models built via Gypsum-DL (Ropp et al., 2019b, 2019a). Different ionization states were obtained by considering pH values between 6.4 and 8.4. The crystallographic structure of HSA (PDB id: 1AO6) was used as a target (Sugio et al., 1999). The system preparation, namely formatting input, protonation, charges computation, and setting of rotatable bonds in the ligand and protein residues, was carried out via Autodock Tools (Morris et al., 2009). Finally, the flexible docking was carried out via Autodock Vina (Trott and Olson, 2010). The well-known and characterized Sudlow site located at Subdomain IIA, referred to as site 1, was not investigated given the FRET results suggest distances incompatible with binding on this binding site. A second site that occurs at domain IIB was then considered, and site residues ARG114, ARG117, ASP183, ARG186, LYS190 and LYS519 were set flexible the docking calculations.

## 3. RESULTS AND DISCUSSION

### 3.1. Mechanism of interactions by fluorescence quenching

Fluorescence quenching is one of the most used techniques for proteins and ligand interactions. This process occurs when the aromatic amino acid residues (Phe, Tyr, Trp, which are responsible for the fluorescence emission of proteins) are suppressed due to the interaction with small molecules. Therefore, HSA fluorescence (the free protein showed λ_em_ = 335 nm when excited at 280 nm) was monitored during DR interactions at three temperatures. Fig.2 shows the HSA fluorescence quenching caused by increasing DR concentrations for the temperatures of 296 K (a), 303 K (b) and 310 K (c). The fluorescence quenching phenomenon can originate from a series of intermolecular processes, such as ground-state complex formation, collisional quenching, molecular rearrangements, excited state reactions, and transfer energy (D. Zhang et al., 2018). In general, the fluorescence quenching mechanism can be classified as dynamic or static. The dynamic mechanism originates from the collision between the fluorophore and the quencher. In contrast, the static mechanism is associated with forming a complex in the excited state that persists in returning to the fundamental state (Lakowicz, 2013).

**Figure 2.**
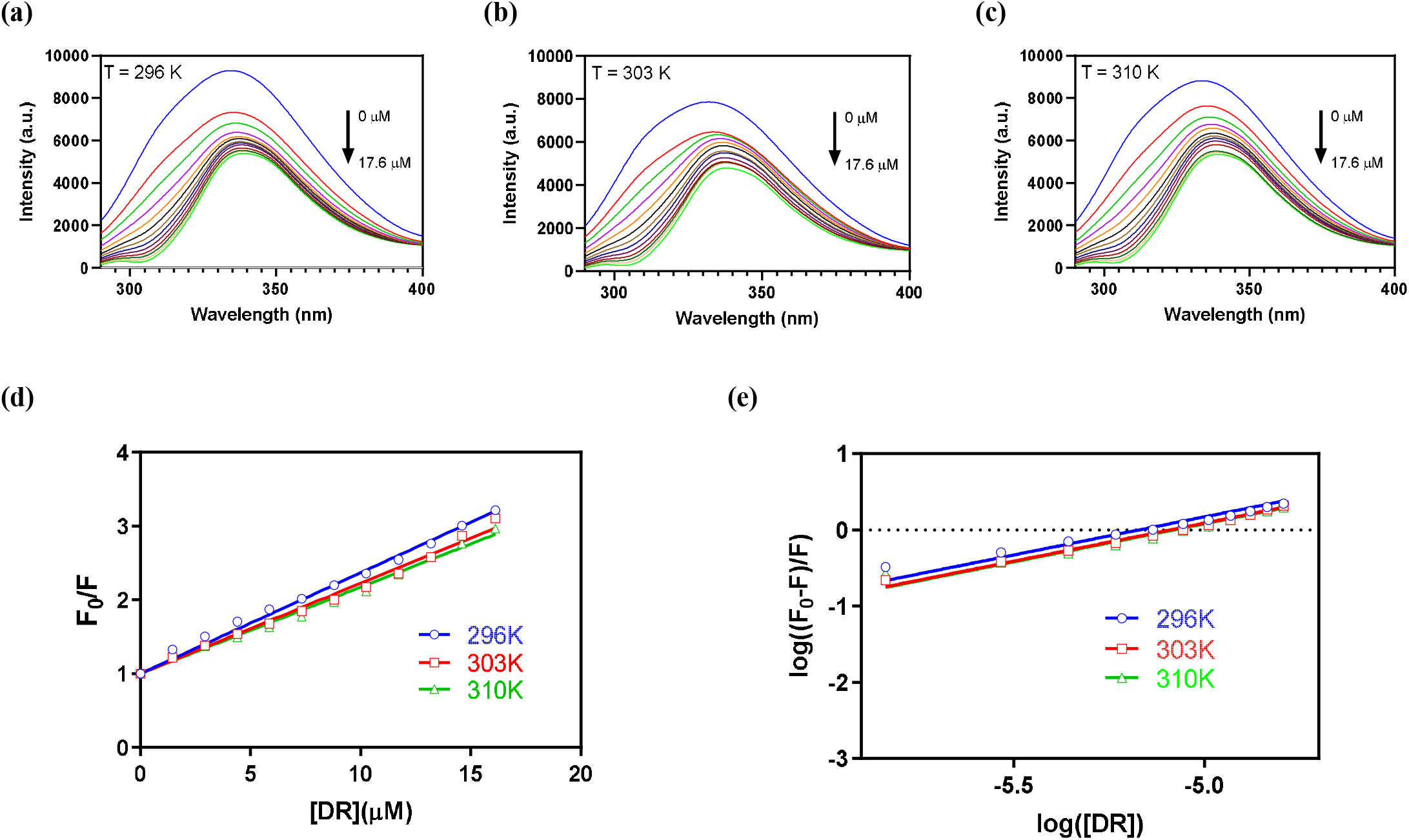
HSA fluorescence quenching spectra at 296 K (a), 303 K (b), and 310 K (c), pH 7.4. Stern-Volmer plots (d) and modified Stern-Volmer plots (e) of HSA-DR interactions.

The Stern-Volmer graphs (Fig. 2d) and their temperature dependence were monitored to detail the HSA quenching mechanism by DR. Static mechanisms are characterized by a K_sv_ decrease, occasioned by increasing temperature since the binding in the formed fluorophore-quencher complex are weakened at high temperatures. In contrast, interactions increase due to increased collisions at higher temperatures for dynamic mechanisms (Fan et al., 2013; Hu et al., 2005). Table 1 shows the calculated Stern-Volmer (K_sv_) constants decrease with increasing temperature, suggesting the static mechanism predominance in HSA-DR interactions. Furthermore, the k_q_ values presented in table 1 are consistent with the static mechanism, considering that the constants were higher than the maximum scattering collisional quenching constant (2×10^10^M^−1^s^−1^) (Alsaif et al., 2020; Fathi et al., 2019).

**Table 1.** Binding parameters of the fluorescence quenching of HSA-DR interactions.

| *Temperature (K)* | **$K_{sv} \pm SD (10^5 M^{-1})$** | **$K_q (10^{13})$** | **$K_b \pm SD (10^5 M^{-1})$** |
| --- | --- | --- | --- |
| 296 | $1.36 \pm 0.02$ | 2.14 | $1.48 \pm 0.01$ |
| 303 | $1.22 \pm 0.02$ | 1.91 | $1.23 \pm 0.01$ |
| 310 | $1.17 \pm 0.01$ | 1.84 | $1.19 \pm 0.01$ |

### 3.2. Binding parameters and number of binding sites

The molecular affinity is a key property to characterize the interaction of xenobiotics with transport proteins, considering that the biodistribution of small molecules in the body and their potential for toxicity can be assessed by this parameter (Poór et al., 2017). From a toxicological point of view, moderate interactions are worrisome due to the high efficiency in the biodistribution of dangerous substances, bringing irreversible damage to human health (Rao et al., 2020). In contrast, weak interactions lead to short half-lives and prevent the transport of the compound by carrier proteins, while very strong interactions reduce the concentration of the free substance in the plasma due to rapid elimination by the body (Chen et al., 2011; de Araújo Motta et al., 2021). The binding constant (K_b_) were obtained from the modified Stern-Volmer graph (Fig. 2 (e)), with K_b_ values in the range of 1.48-1.19×10^5^ M^−1^. Thus, K_b_ values represent moderate interactions, with similar values already reported in previous studies of dye-albumin interaction (Ksenofontov et al., 2019; Naveenraj et al., 2018, 2013).

### 3.3. Mode of binding and thermodynamic parameters

Thermodynamic parameters are used to identify the guiding forces involved in the HSA-DR complex. Fig. 3a shows the ΔG° vs. temperature plots used to determined ΔS° and ΔH° (according to Eq. 4 and 5) and Fig. 3(b) displays the determined thermodynamic parameters: ΔG ° = −29.26 kJ.mol^− 1^ (296 K), −29.49 kJ.mol^−1^ (303 K) and −30.10 kJ.mol^−1^ (310 K); ΔS° = 59.93 J.mol^−1^ and ΔH ° = −11.46 kJ.mol^−1^. The interaction HSA-DR is spontaneous for all considered temperatures since ΔG° values are negative. In addition, bearing in mind that the values of ΔS° > 0 and ΔH° < 0, it suggests that electrostatic interactions govern the interaction (Ross and Subramanian, 1981).

**Figure 3.**
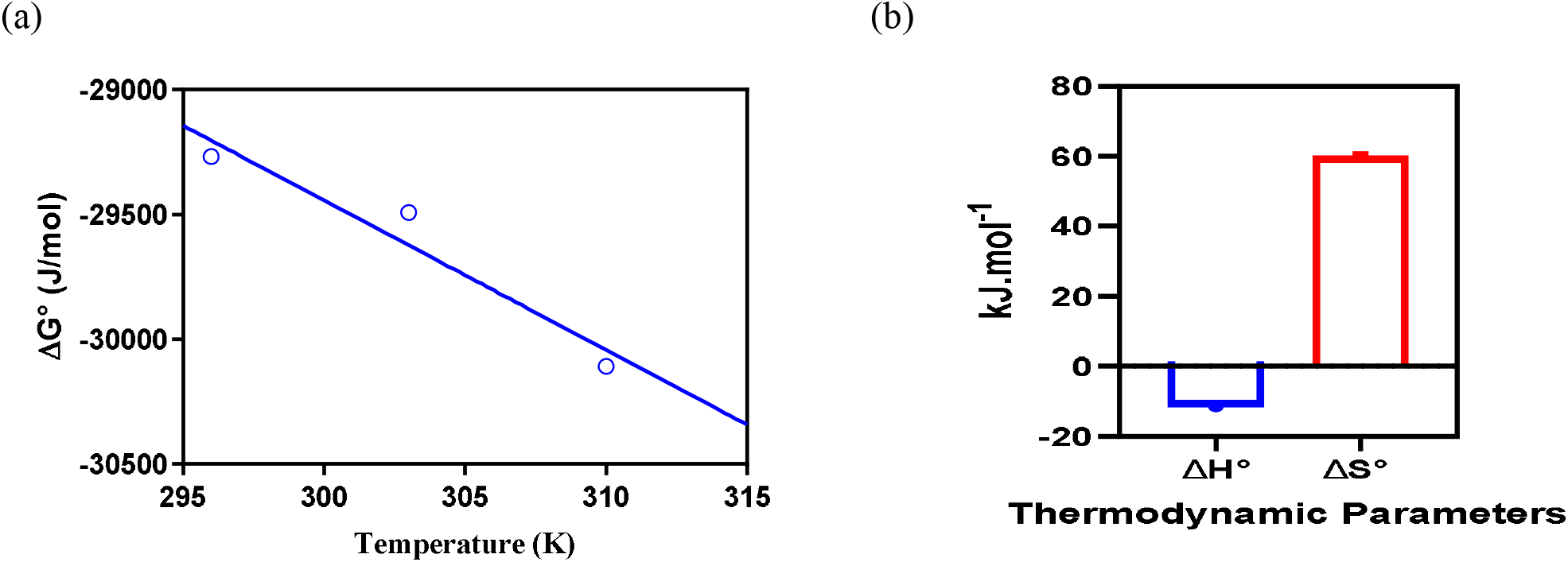
Van’t Hoff chart with ΔG vs. Temperature (a) and thermodynamic parameters (b) of HSA-DR interactions.

### 3.4. Determination of the binding distances and energy transfer parameters

Interaction processes between fluorophores of biomacromolecules (donors) and small molecules (acceptors) may involve transferring non-radioactive energy between them. Here, The FRET studies were carried out according to Forster’s theory to characterize this energy transfer. FRET occurs when the acceptor (DR) absorption overlaps with the donor (HSA Trp-214 residue) emission spectrum. In addition, the acceptor-donor distance must be less than 10 nm (Chen et al., 2020). Fig. 4 represents the overlap of the HSA emission spectrum with the DR absorption spectrum, demonstrating the possible occurrence of FRET between such species. The parameters obtained through the FRET study are shown in table 2. Through equation 6, it was possible to obtain R_0_ = 3.87 nm and r = 5.52 nm. Note that such values fit perfectly in theory proposed by Forster, with values <10 nm and satisfying the 0.5R_0_ <r <1.5R_0_ dependency, suggesting the presence of non-radioactive energy transfer in the HSA-DR system.

**Figure 4.**
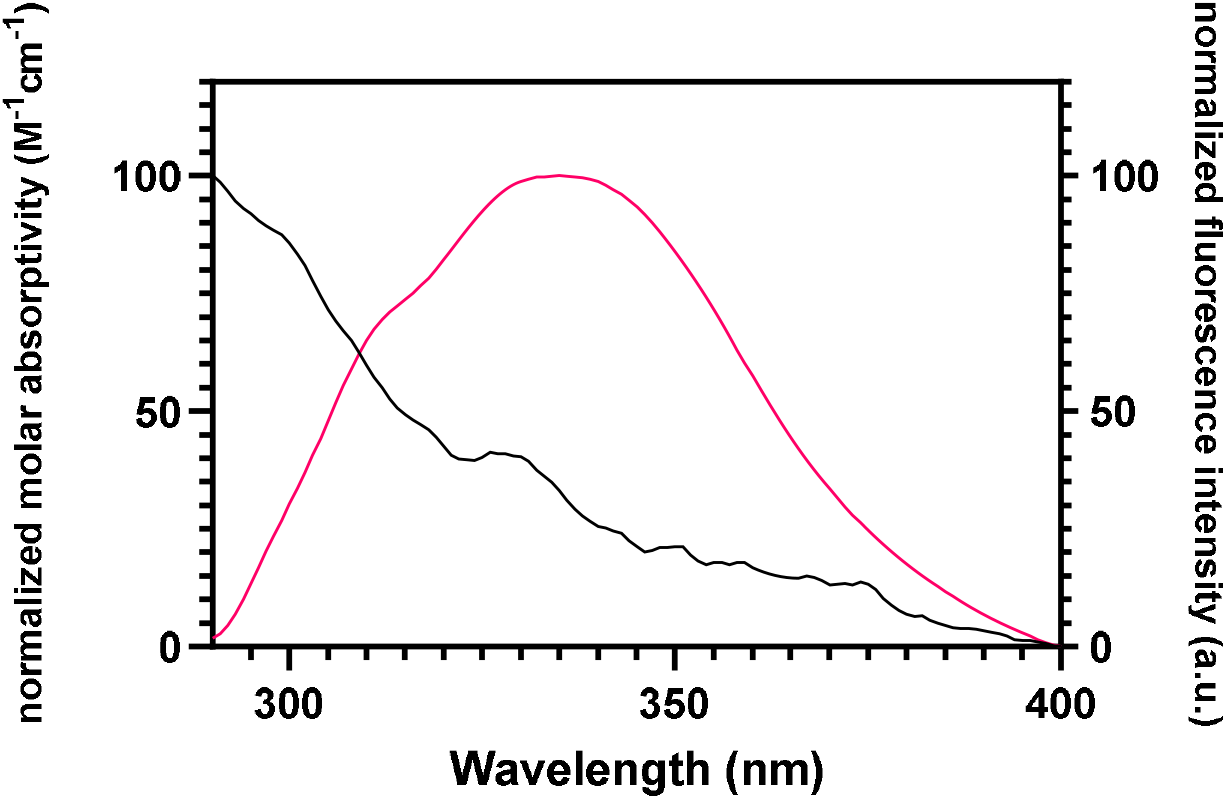
Overlapping of the fluorescence emission spectrum of HSA with the absorption spectrum of DR at 298 K, pH 7.4. [HSA] = [DR] = 0.6 μM.

**Table 2.**
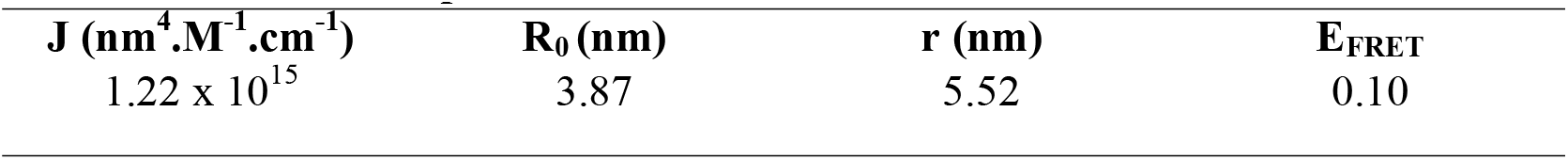
FRET parameters obtained from the HSA-DR interactions.

### 3.5. Analyzing the microenvironment of fluorophores by synchronous fluorescence

Synchronous fluorescence spectroscopy can provide information regarding the neighborhood around HSA aromatic residues (Tyr and Trp) when interacting with DR (Shaghaghi et al., 2019). This technique is very sensitive to changes in polarity of the protein fluorophores, being performed through the simultaneous measurement of excitation and emission wavelengths, under the condition of Δλ = λ_emission_−λ_excitation_, with Δλ = 60 nm for Trp residues and Δλ = 15 nm for the study of Tyr residues (Bagalkoti et al., 2019). The synchronous fluorescence spectra for Δλ = 15 nm and Δλ = 60 nm are displayed in Fig. 5. The percentage of the synchronous fluorescence quenching ratio (RSFQ%) of the Tyr and Trp residues was calculated using equation 9. As seen in Fig. 5(c), at the maximum concentration of added DR (17.6 μM), there is a reduction in the fluorescence of Tyr residues by 84.7% and Trp by 20.82%. Thus, the RSFQ values of the Tyr residues are much higher when compared to the values of the Trp residue, demonstrating a greater contribution of the Tyr residues in the HSA-DR interaction process. In addition, such interactions cause a redshift for the Tyr residues (300 to 301 nm) and a blue shift for the Trp residues (340 to 339 nm), proving an increase in polarity decrease in hydrophobicity around the Tyr residues, in contrast to the Trp residues. Additionally, it can be noticed that there is no formation of the HSA-DR complex at binding site I, given the weak interaction with the only tryptophan residue (Trp-214) located in the subdomain IIA (Rao et al., 2020).

**Figure 5.**
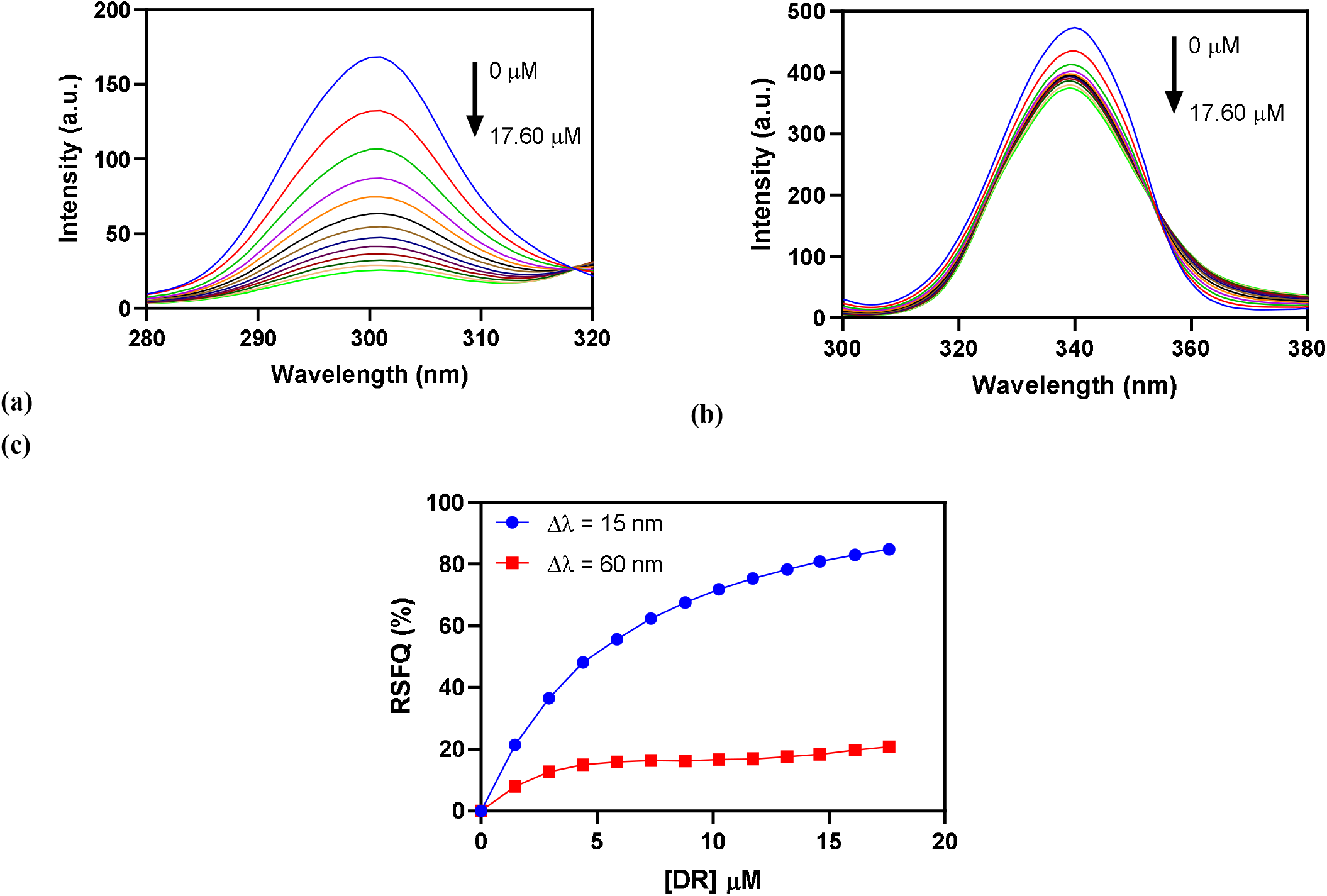
Synchronous fluorescence spectra of HSA-DR systems with Δλ = 15 nm (a) and with Δλ = 60 nm. RSFQ (%) values in increasing concentrations of DR (c). [HSA] = 0.6 μM; [DR] = 0 – 17.60 μM.

### 3.6. Modulation of the HSA UV-Vis spectra by DR

The UV-Vis technique is an excellent tool to verify the formation of a protein-ligand complex in the fundamental state and changes in the polypeptidic structure of the protein (Bratty, 2020; Santos et al., 2018). The formation of this complex is confirmed by changes in the characteristic wavelengths of the UV-Vis spectrum of the protein when it comes into contact with the test substance, such as changes in the absorption intensity and peak displacements (Ahmad et al., 2020; Đukić et al., 2020). Fig. 6 shows the UV-Vis spectra of HSA after successive additions of DR (from 0 to 17.6 μM). The interference of DR in the spectrum of HSA was eliminated using the spectrum of solution corresponding to the concentrations of DR.

**Figure 6.**
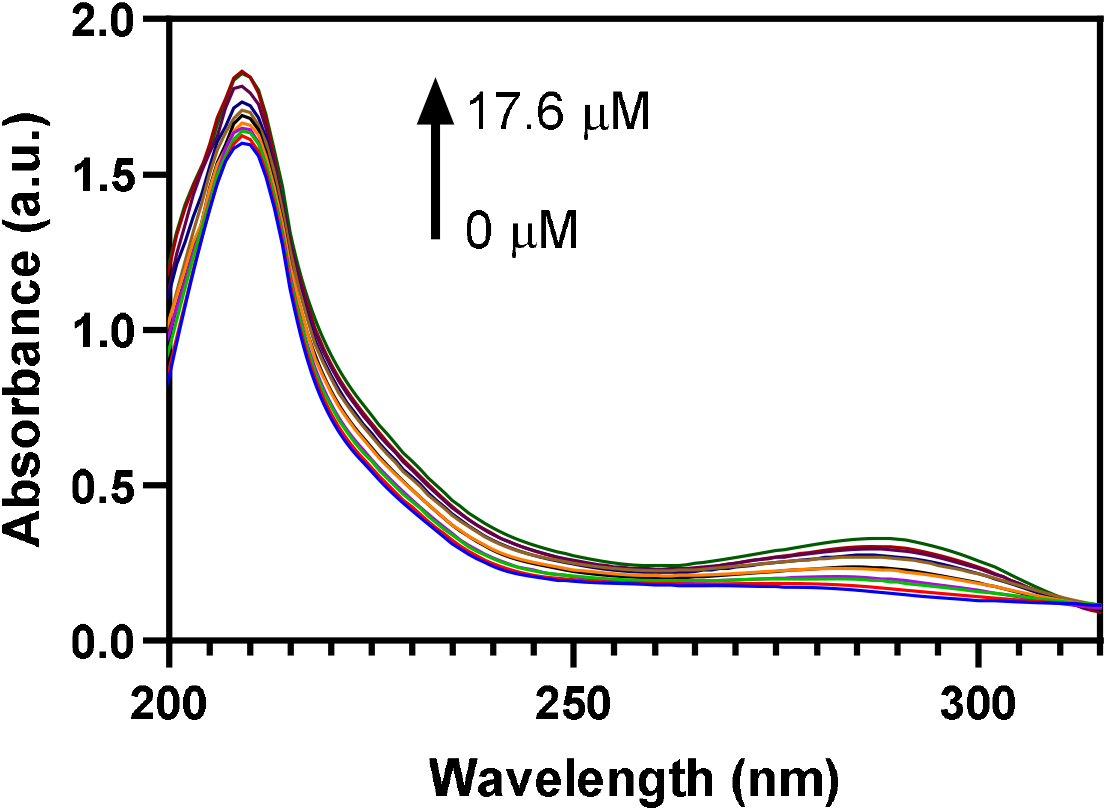
Absorption spectrum of HSA (a) with increasing concentrations of DR at 298 K, pH 7.4. [HSA] = 0.6 μM; [DR] = 0 to 8.8 μM.

The HSA spectra demonstrate two absorption peaks related to the structure of the protein backbone and the π-π * transitions of the aromatic amino acid residues Phe, Trp and Tyr (Vidhyapriya et al., 2019), at 209 and 277 nm, respectively. An increase in HSA absorption intensity was observed at the peak of 209 nm by additions of DR. On the other hand, the peak at 277 nm (Fig.6) shows an increase in the absorption intensity concomitant with the increase in the DR concentration and a redshift of Δλ = 9 nm. These results suggest that the HSA-DR interaction results in changes in protein conformation and confirm the changes in the microenvironment of the fluorophore observed in synchronous fluorescence (Shahabadi and Zendehcheshm, 2020).

### 3.7. Estimation of HSA thiol group

The reactivity of the HSA-SH groups and their effect on exogenous substances is a parameter of great relevance. For example, the thiol group of cys-34 has a great tendency to form covalent adducts with certain xenobiotics due to its high nucleophilicity (Ndreu et al., 2020). Besides, the formation of covalently binding adducts can damage the structure and function of the HSA (Shu et al., 2019), promoting the development of health problems, such as contact allergies (Chipinda et al., 2011), cancer (Nunes et al., 2019) and cardiovascular diseases (Xiao et al., 2018).

For evaluating this parameter, the Ellman test was performed, which is based on the reaction between the DTNB and the SH-HSA, generating the TNB^−^ product identified by absorption at 412 nm. Fig. 7 represents the number of free thiol groups in the studied system. A decrease in SH groups free of HSA was observed according to the increase in DR, reaching a reduction of 31.11% in the reactive thio groups in the highest concentration of DR. This can mean the occurrence of two situations: (1) part of the SH groups did not show reactivity due to the formation of the DR-Cys34-HSA adduct; (2) DR is inducing a change in the structure of the HSA so that the Cys34 residue becomes inaccessible to react with the DTNB.

**Figure 7.**
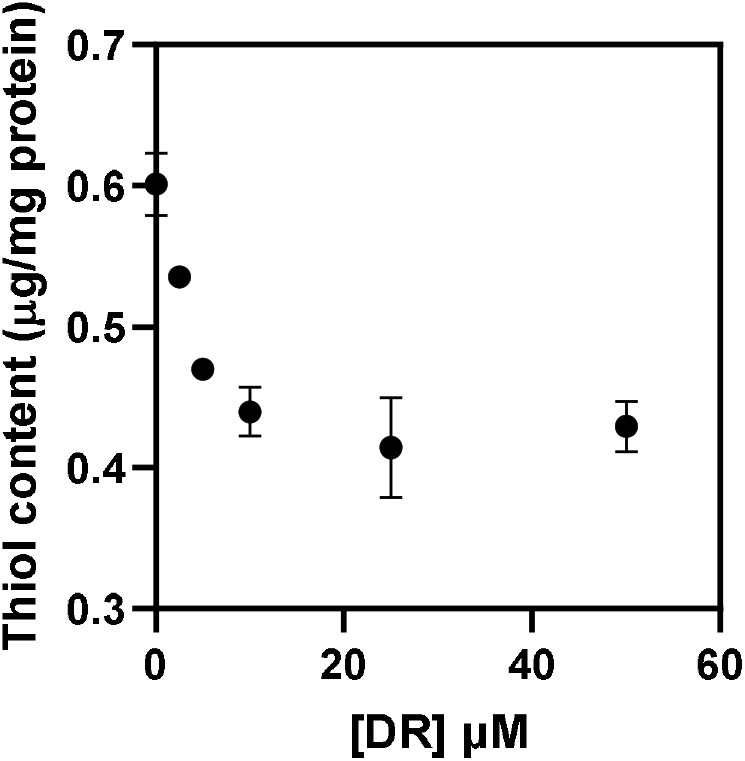
Amount of free thiol group in HSA treated at various concentrations of DR.

### 3.8. Rayleigh light scattering (RLS) spectroscopy measurements

RLS measurements were performed to verify changes in protein particle volume after interaction with DR. It is widely known that the intensity of the RLS increases proportionally with the volume of the dispersion particles (Pasternack and Collings, 1995). Fig. 8 shows the HSA RLS intensities at 350 nm in the absence and presence of increasing concentrations of DR. The intensity of the RLS decreased with the increase in DR in the system, suggesting a reduction in the size of the HSA particles. A plausible explanation for this result is the destruction of the solvent shell present on the protein’s surface, leading to greater dispersion of the HSA. Another possible situation is the collapse of the protein when interacting with DR, causing HSA contraction due to conformational changes, leading to the reduction or loss of biomolecule function (Wang et al., 2018; Xu et al., 2019).

**Figure 8.**
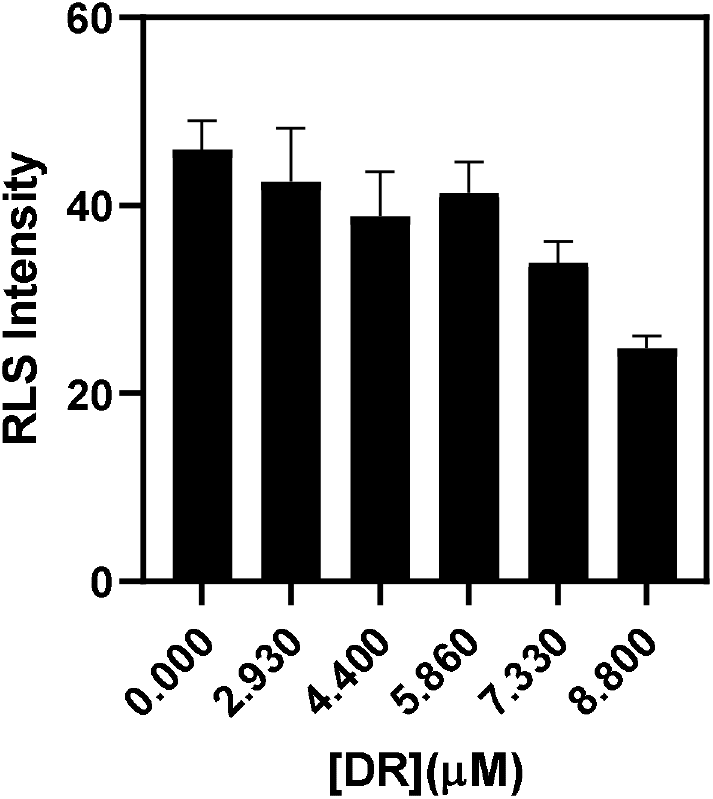
RLS intensities at λ_ex_ = λ_em_ = 350 nm represented in a bar diagram.

### 3.9. Molecular Docking

The FRET results suggest a 5.52 nm mean distance between the DR molecule and W214 residue of HSA. W214 is located at the well-known and characterized Sudlow site of HSA. Therefore, it is unlikely that DR has this site as its main binding site. For that reason, a second site located at domain IIB was investigated. Docking analyses estimate an affinity of 10.8 kcal/mol for the best pose obtained (Fig. 9), and the average estimate for the top 10 best poses is 10.5 kcal/mol. The best pose suggests that site 2 can provide hydrogen bonds via R114, H146, Y161 and R186, and hydrophobic contacts via L115, I142, K190, E425, P421 and I523. The average distance computed from the distance of the pairwise atoms between the poses obtained and residue W127 is 2.7 nm (σ = 0.35), which is closer to the 5.52 nm. The flexible docking of DR suggests that this cavity can provide a stable positioning for DR. The interactions include histidine and tyrosines, which are compatible with the fluorescence reduction observed for the HSA-DR system. This site also seems like a plausible site for disrupting the structural stability of domain IIB. The domain IIB presents 4 helices and a long loop that connects it to domain IA (Sugio et al., 1999). It is possible that a ligand binding that could disrupt the loop structure, especially if there is available space for repositioning the loop, could promote an increase in the general flexibility of the domain. The Van’t Hoff charts discussed points that this is the case for HSA-DR binding (Fig. 3). There is evidence for the protein conformational entropy change to be the main driving force of the total binding entropy change for general protein-ligand binding events (Tzeng and Kalodimos, 2012; Verteramo et al., 2019). Therefore, this hypothesis would follow the experimental results and point to future research directions necessary to elucidate the atomistic details of HSA-DR binding.

**Figure 9.**
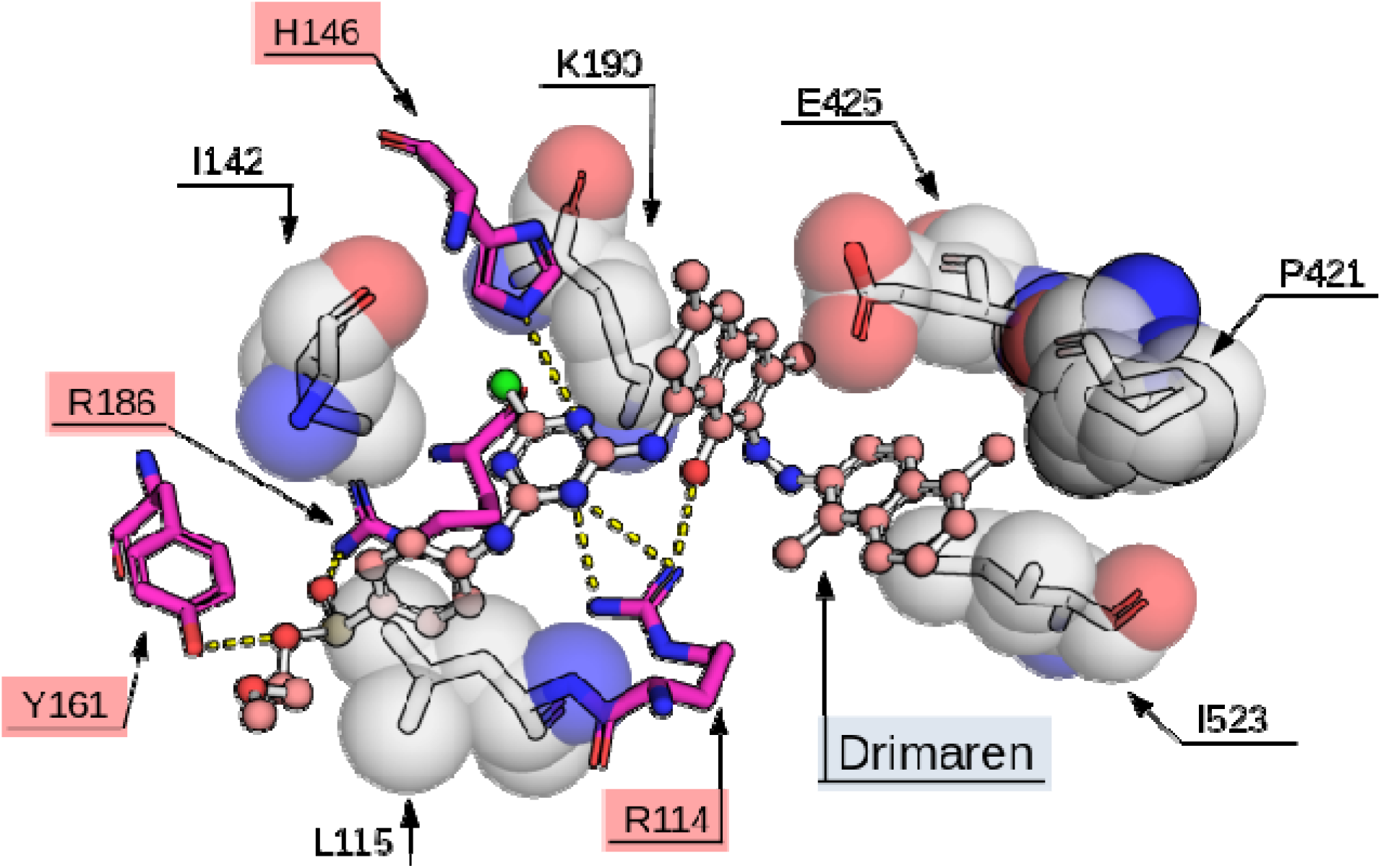
DR best docking poses at HSA domain IIB site. Spheres represent the Van der Walls radius of residues atoms involved in ligand hydrophobic contact and dashed lines represents hydrogen bonds.

## 4. CONCLUSION

We systematically investigated the molecular interaction between DR and HSA by multispectroscopic and molecular docking approaches. The fluorescence spectroscopy results indicated that DR could cause a static quench to the intrinsic fluorescence of HSA. Thus, the binding interactions between DR and HSA formed complexes that were stabilized mainly by electrostatic interactions. Synchronous fluorescence and UV-Vis studies proved that some microenvironmental and conformational changes in HSA were induced by their binding interactions, indicating the potential toxicity of DR to cause the damage of secondary structure and the disturbance of normal biological functions of HSA. Ellman’s assay demonstrates a possible formation of HSA-DR adduct or structural change of HSA when interacting with DR. RLS shows conformational damage associated with HSA interactions. Moreover, molecular docking suggests the binding site of DR was located in subdomain IIB HSA. Overall, these studies provide clear insight into the molecular interaction between DR and proteins, which may prove useful for evaluating the toxicity and effects of DR in biological systems.

## Acknowledgments

This work was supported by the Foundation for Science and Technology of Pernambuco FACEPE (grant number AMD-0096-1.06/19). Additionally, the authors are grateful to the Laboratory of Organic Compounds in Coastal and Marine Ecosystems at the Federal University of Pernambuco (OrganoMAR - UFPE) and the Laboratory of Enzymology (LABENZ - UFPE) for providing infrastructure to carry out the fluorescence experiments.

